# Xpo7 negatively regulates Hedgehog signaling by exporting Gli2 from the nucleus

**DOI:** 10.1101/2020.01.31.928408

**Authors:** Łukasz Markiewicz, Tomasz Uśpieński, Sylwia M. Niedziółka, Paweł Niewiadomski

**Affiliations:** Laboratory of Molecular and Cellular Signaling, Centre of New Technologies, University of Warsaw, Banacha 2c, 02-097 Warsaw, Poland

## Abstract

Dynamic bidirectional transport between the nucleus and the cytoplasm is critical for the regulation of many transcription factors, whose levels inside the nucleus must be tightly controlled. Efficient shuttling across the nuclear membrane is especially crucial with regard to the Hedgehog (Hh) pathway, where the transcriptional signal depends on the fine balance between the amounts of Gli protein activator and repressor forms in the nucleus. The nuclear export machinery prevents the unchecked nuclear accumulation of Gli proteins, but the mechanistic insight into this process is limited. We show that the atypical exportin Xpo7 functions as a major nuclear export receptor that actively excludes Gli2 from the nucleus and controls the outcome of Hh signaling. We show that Xpo7 interacts with several domains of Gli2 and that this interaction is dependent on SuFu, a key negative regulator of Hh signaling. Our data pave the way for a more complete understanding of the nuclear shuttling of Gli proteins and the regulation of their transcriptional activity.

## Introduction

The Hedgehog pathway (Hh) is an evolutionarily conserved signaling cascade that regulates embryonic patterning and organ morphogenesis by orchestrating cell proliferation, differentiation, and migration (1–4). Aberrations of Hh signaling result in birth defects, such as polydactyly or cyclopia, and cancer. Hyperactivation of this pathway often occurs in cancer and drives aberrant cancer cell proliferation and self-renewal (1, 5).

In the Hh cascade, the concentration of the ligand is translated via the receptor Patched (Ptch) and a transmembrane domain protein Smoothened (Smo) into the activity of the bifunctional Gli transcription factors, which are the transcriptional effectors of the Hh pathway (6). In mammals, they are represented by three proteins: Gli1, Gli2, and Gli3. Gli1 acts principally as a transcriptional activator, whereas Gli2 and Gli3 display both activator and repressor functions. When the pathway is off, Gli2 and Gli3 are partially truncated on the C terminus. This cleavage removes their transcriptional activation domain and converts them into repressors of Hh target genes. When Hh ligand binds to the Ptch receptor, activated Smo blocks the proteolytic truncation of Gli proteins. Full-length Gli proteins are posttranslationally modified and then translocate into the nucleus. In there, they recruit transcriptional co-activators and induce the expression of Hh target genes, including *Gli1* (7, 8). The expression of target genes is precisely regulated by the balance between Gli protein activator and repressor forms inside the nucleus. This balance is achieved via nucleocytoplasmic shuttling of Gli proteins, a process that up till now was only partially understood.

The nucleocytoplasmic exchange of transcription factors through nuclear pore complexes (NPCs) is a key regulatory mechanism in many signaling pathways (9). Specific sequence motifs known as nuclear localization (NLS) and nuclear export signals (NES) are recognized by karyopherin nuclear transport receptors: respectively importins and exportins. These transport receptors ferry cargos across NPCs along the gradient of different forms of the small GTPase Ran (10). Importins bind cargos in the cytoplasm where Ran is present primarily in its GDP-bound form. They release their target proteins in the presence of RanGTP in the nucleus (11, 12). Conversely, the binding of exportins to their substrates is strong in the presence of RanGTP and cargos are released when RanGDP is present (13). Of the 8 mammalian exportins, the best studied by far is Xpo1/Crm1, a broad-specificity transport protein with well-established recognition sequences. Among its other cargos, Xpo1 was shown to participate in the nucleocytoplasmic transport of Gli1, but Gli2 cytoplasmic export appears to be Xpo1-independent (14). Full length activator forms of Gli2 and Gli3 are mostly excluded from the nucleus in untreated cells, but they do localize in the nuclear fraction to some extent (7, 15). This suggests that either some baseline active import or slow diffusion of Gli2/3 proteins into the nucleus occurs even in cells not exposed to the Hh ligand. We hypothesized that one or more of the exportins counteracts this inward flux of Gli2/3 through the nuclear pore to ensure that Gli proteins do not accumulate in nuclei in an unrestrained manner. We reasoned that the loss of function of this exportin would result in unchecked buildup of Gli2/3 activators in the nucleus and the activation of target gene transcription even in the absence of the ligand. Data from a recent CRISPR/Cas9-based functional screen for regulators of Hh signaling suggested that the atypical exportin Xpo7 may play such a role (16), and we chose this exportin for further analysis.

RanBP16/Xpo7 was initially characterized as an exportin based on its ability to directly interact with RanGTP (17), but its role in physiological processes has been poorly studied (17–19). It is strongly induced in erythropoiesis, and its knockdown interferes with nuclear condensation, halts the loss of histones from nuclei, and blocks enucleation in erythroblasts (20). Recent work identified over 200 substrates for Xpo7 using affinity chromatography and mass spectrometry (18). Unlike for Xpo1, Xpo7 substrates do not carry short linear NES motifs and are likely recognized by Xpo7 through more diffuse signals, making the task of characterizing Xpo7-interacting protein domains challenging. NESs for Xpo7 generally contain basic residues and folded motifs, although Xpo7 has also been shown to facilitate the export of classical NES-containing cargoes, suggesting that the substrate specificities of Xpo1 and Xpo7 may partially overlap (18).

Here we show that Xpo7 functions as a nuclear export receptor that excludes Gli2 from the nucleus and plays a role in controlling the outcome of Hh signaling. We also find that the binding to Xpo7 is mediated by the N-terminal domain and the A1 domain of Gli2. Moreover, we demonstrate that the depletion of Xpo7 results in changes in Hh pathway activation. This establishes Xpo7 as a Gli protein transporter and paves the way for a more complete understanding of the function of nuclear import/export pathways in Hh signaling.

## Materials and methods

### Constructs and cell lines

Gli and Xpo7 constructs described in this study were made from mouse cDNA. Coding sequences or their fragments were amplified by PCR and cloned into the pENTR2B (Life Technologies) vector by Gibson assembly (NEBuilder® HiFi DNA Assembly Master Mix; NEB). Gli protein constructs were tagged with the N-terminal 3xHA tag in tandem with the TEV protease target sequence. Constructs were then shuttled into the pEF/FRT/V5-DEST (Life Technologies) vector using Gateway cloning in frame with the C-terminal V5 tag (Gateway LR Clonase II mix; Life Technologies). The Gli2 P1-6A mutant was previously described (15). RanQ69L mutant was a kind gift from professor Ian G. Macara and previously described (21). RanQ59L and mouse SuFu were subcloned into pcDNA3.1 in frame with an N-terminal 2xFLAG-tag.

Stable cell lines expressing low levels of HA-tagged Gli2 and Gli3 variants were generated using the Flp-In system according to the manufacturer’s recommendations (Life Technologies) (22). Briefly, NIH/3T3 Flp-In cells were co-transfected with pOG44 and the pEF5/FRT/V5-DEST vector containing the construct of interest. After 2 days the cells were reseeded at low density and the culture media was supplemented with hygromycin for stable integrant selection. Stable cell lines were reselected with hygromycin on every other passage to preserve selection pressure and prevent silencing of the transgene.

For transient transfections of cells JetPrime reagent (Polyplus) was used according to the manufacturers protocol. For siRNA transfection Lipofectamine RNAiMAX Reagent (Thermo) was used. Each siRNA (ON-TARGETplus siRNA Reagents, Horizon) was introduced at 40 pmol/well for 48h.

### Cell culture

HEK-293T cells (ATCC), and NIH/3T3 Flp-In cells (Life Technologies), including derivative stable clones were cultured in media composed of DMEM (high glucose), 10% fetal bovine serum (FBS; Thermo Fisher Scientific), 1x GlutaMAX, 1x non-essential amino acids, 1x sodium pyruvate, 1x penicillin/streptomycin (all from Life Technologies). Prior to harvesting, the cells were serum-starved in the same media but containing 0.5% FBS for 24 hours with or without SAG (400 nM; Enzo) depending on the experiment.

### CRISPR/Cas9 KO generation

The gRNA sequence: 5’-TTAAAGAGTAACTACCAACT-3’ targeting the mouse *Xpo7* gene was cloned into the PX459 plasmid (Addgene #62988, (23)). 8×10^5^ of NIH/3T3 Flp-In cells were seeded in a 24-well plate 2h prior to the transfection. Cells were transfected with 500 ng PX459-sgXpo7 per well using the PTG1 reagent (24). 48h post-transfection the cells were selected with 2ug/mL puromycine for 2 days. To isolate clones from single cells, the cells were seeded at a density of 0.5 cells per well on a 96-well plate. 4-5 days later cells were expanded from wells where a single colony was present.

### Luciferase Reporter Assay

The Hedgehog reporter luciferase assay was performed as described before (15). Cells were transiently transfected with a mixture (4:1) of a firefly luciferase reporter driven by a Gli-responsive promoter and a constitutive Renilla luciferase reporter on a 48-well plate (25). Cells were grown to confluence and switched to a medium containing 0.5% FBS. Dual luciferase assay was performed using the Dual-Luciferase Reporter Assay System (Promega) on the Synergy H1 plate reader (Biotek).

### Western Blot Analysis and Co-immunoprecipitation

HEK-293T cells were harvested 24h post-transfection. For the production of whole-cell lysates, cells were lysed in 300 μl of lysis buffer (50 mM Tris at pH 7.4, 1% NP-40 [v/v], 150 mM NaCl, 1mM GTP, 10mM MgCl_2_, a protease inhibitor cocktail [1x EDTA-free protease inhibitors, Sigma], 10mM NaF, 1mM Na_3_VO_4_, 2μM Bortezomib) at 4°C. 60 μl of lysates was saved as input and the rest was subject to immunoprecipitation.

For immunoprecipitation, rat anti-HA antibodies (Roche) were coupled covalently to protein G-coated magnetic beads (Dynal) using dimethyl pimelimidate. Anti-HA-coupled beads were added to the clarified lysate. After 1.5h binding at 4°C, the beads were washed 3x 5min at 4°C with lysis buffer and eluted with 2x SDS sample buffer at 65°C for 30 min.

Western blot detection was performed by enhanced chemiluminescence using HRP-coupled secondary antibodies and the Clarity or Clarity Max substrates (Bio-Rad). The signal was digitized using Amersham 600RGB imager as 16-bit grayscale TIFF files. Quantitative analysis of band intensities was performed with ImageJ.

### qRT-PCR

RNA was isolated using the Trizol method according to the manufacturer’s instructions. Reverse transcription was performed using the High-Capacity cDNA Reverse Transcription Kit (Thermo). Real-Time qRT-PCR was performed using the Real-Time 2xHS-PCR Master Mix Sybr B (A&A Biotechnology) on a LightCycler 480 II qPCR System (Roche) with custom-designed primers for *Gli1* (Fwd: 5’-CCAAGCCAACTTTATGTCAGGG-3’ and Rev: 5’-AGCCCGCTTCTTTGTTAATTTGA-3’), *Xpo7* (Fwd: 5’-GAGCAAAATGGCGGATCATG-3’ and Rev: 5’-AGTCGAGTGGTTGTATCTGTTG-3’), and *Gapdh* (Fwd: 5’ - GGCCTTCCGTGTTCCTAC-3’ and Rev: 5’-TGTCATCATACTTGGCAGGTT-3’). Transcript levels relative to *Gapdh* were calculated using the ΔΔCt method.

### Immunofluorescence and microscopy

Cells were cultured on glass coverslips in a 24-well plate. Cells were fixed in 4% [w/v] paraformaldehyde in PBS for 10 min at room temperature (RT) and washed in PBS (3 × 5 min). Fixed cells were incubated with permeabilization buffer (0.3% Triton X-100 in PBS) for 15 min at RT. Then, cells were washed in PBS (3 × 5 min) and blocked in blocking buffer (1% [v/v] normal donkey serum, 10 mg/ml [w/v] bovine serum albumin, 0.1% [v/v]Triton X-100 in PBS) for 1h at RT. Primary antibodies were diluted in blocking buffer and used to stain cells overnight at 4°C. After washing (3 × 5 min) in PBS containing 0.1% [v/v] Triton X-100), secondary antibodies were added in blocking buffer and incubated for 1 h at RT. The cells were washed as above and mounted with DAPI-containing anti-fade medium (ProLong Diamond, Thermo). Microscopy was performed on an inverted Olympus IX-73 fluorescent microscope equipped with a 63x oil objective and the Photometrics Evolve 512 Delta camera.

For the quantitative analysis of fluorescence intensities, all images were obtained with identical gain, offset, and mercury lamp power settings. Nuclei and cytoplasm were manually annotated and fluorescent intensities were measured with the help of an ImageJ script. To quantify the fraction of Gli or truncated construct in the nucleus or cytoplasm, we calculated the log_10_ values of the ratios of intensities of the fluorescent signal in the nuclear and cytoplasmic fractions in each cell. When using shRNA transient transfection, we co-transfected a tdTomato expression plasmid to distinguish cells transfected with Xpo7 shRNA from untransfected cells. Only cells with clear red fluorescence were used for downstream analysis.

### Antibodies

The following antibodies were used for immunoprecipitation, western blot, and immunofluorescence: HA (high affinity) no. 11867423001 (Roche); DYKDDDDK (FLAG) no. 635691(Clontech); Exportin 7 (A-11) no. sc-390025 (Santa Cruz); V5 tag no. ab27671 (Abcam); Beta-actin no. A1978 (Sigma); α-tubulin no. T6199 (Sigma). The anti-Sufu antibody was a kind gift from R. Rohatgi and was used previously (7).

### Data analysis

All statistical tests were performed using R/RStudio or Excel. Student’s t-test was used for assessing statistical significance. For bar graphs, bar heights are mean measurements and error bars represent standard deviation of n=3 measurements, unless otherwise indicated. For box plots, boxes show median, upper and lower quartile values and whiskers show maximum and minimum values, excluding outliers (which are shown separately as circles).

## Results

### 1. Xpo7 depletion affects Hh pathway activity

To determine the role of different components of the nuclear import-export machinery in Hh signaling we reanalyzed the data from a recent CRISPR-Cas9-based genome-wide screen of Hh pathway regulators by Pusapati et al. (16). Based on the phenotype of knockout cells, Xpo7 was classified as a negative regulator of Hh signaling, scoring as one of the top negative regulators under conditions of low pathway activity (cells untreated with Hh agonist - “No Sonic Hedgehog” or treated with suboptimal doses of Sonic Hh - “Low Sonic Hedgehog”). Xpo7 scored higher as a negative Hh modulator than any other exportin, with Xpo1 coming second (Fig. 1A). This data suggests a strong involvement of Xpo7 in Hh pathway regulation. Interestingly, Xpo7 no longer acted as a negative regulator of Hh signaling in the presence of near saturating concentrations of Sonic Hh (Fig. 1A; “High Sonic Hedgehog”).

**Fig.1.**
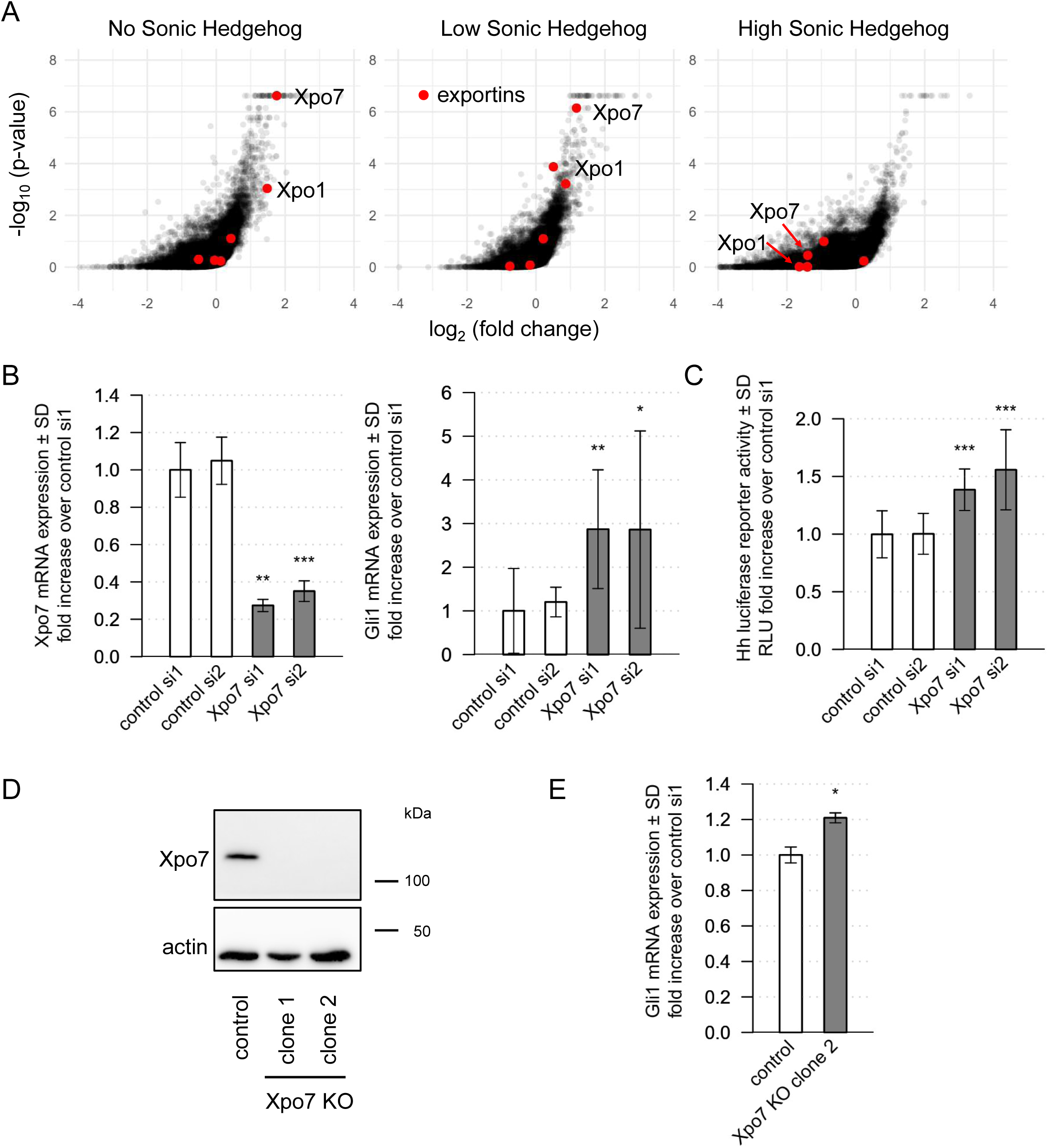
Xpo7 depletion affects Hh pathway activity. **A)** Volcano plots based on CRISPR/Cas9 screens from Pusapati et al., 2018 (16). Cells were transduced with a genome-wide pool of sgRNAs, and gene indel mutations were induced by Cas9. The cells were left untreated (“No Sonic Hedgehog”) or treated with low or near-saturating doses of Sonic hedgehog. Cells with high activity of the Hh pathway were isolated by FACS and sequenced to identify enriched/depleted sgRNAs. The supplemental data from (16) were reanalyzed using a custom R script. Each dot corresponds to one gene targeted by the library, with exportins marked in red. For each gene, the x-axis represents the mean enrichment or depletion of its corresponding sgRNAs in the FACS-sorted population, and the y-axis represents the corresponding uncorrected p-value (16). Genes located towards the top right corner of each graph correspond to negative regulators of Hh signaling **B)** NIH/3T3 cells were treated with two different control siRNAs and two different *Xpo7* siRNAs and mRNA levels for *Xpo7* and *Gli1* were measured by qRT-PCR. *Xpo7* KD increases the activity of the Hh pathway, as assessed by *Gli1* expression **C)** Activity of the Hh luciferase reporter was measured in cells treated with *Xpo7* siRNAs or non-targeting control siRNAs. **D)** NIH/3T3 Flp-In cell lines with CRISPR/Cas9-mediated knockout of *Xpo7* do not have detectable levels of Xpo7 protein. **E)** The expression of the Hh target gene *Gli1* is induced in *Xpo7* knockout cells compared to the unmutated parental NIH/3T3 line (n=3; *** p<0.001; ** p<0.01;* p<0.05).

To directly assess the functional consequences of Xpo7 depletion on Gli-mediated transcriptional activity, we introduced siRNA targeting *Xpo7* into NIH/3T3 fibroblasts. The siRNA-mediated knockdown (KD) resulted in a 70-80% decrease in the level of *Xpo7* mRNA (Fig. 1B; left panel). *Xpo7* KD cells displayed an over 2-fold increase in the expression of *Gli1*, the canonical Hh target gene, compared to control cells (Fig. 1B; right panel), suggesting that the Hh-dependent transcription is induced by *Xpo7* loss. Similarly, the activity of a Gli-dependent luciferase reporter was increased by *Xpo7* KD (Fig. 1C).

To ensure that the effects of *Xpo7* KD were specific, we created *Xpo7* mutant knockout (KO) stable cell lines using CRISPR-Cas9. Two single cell-derived clones of *Xpo7* KO cells showed undetectable levels of Xpo7 protein (Fig. 1D). Similar to *Xpo7* KD cells, *Gli1* expression was increased in *Xpo7* KO cells compared to unmutated controls (Fig. 1E).

### 2. Xpo7 interacts with Gli2 and Gli3 via multiple domains

Given the established function of Xpo7 in nuclear export, we hypothesized that Xpo7 can influence the outcome of Hh signaling by facilitating Gli protein transport out of the nucleus. If that is the case, we expected that Xpo7 would bind to Gli proteins in a RanGTP-dependent manner. Indeed, HA-tagged Gli2 interacts with endogenous Xpo7 in a stable NIH/3T3 Flp-In cell line (Fig. 2A). The interaction is independent of Hh pathway activity – it is present both in untreated cells and in cells treated with the Smo agonist SAG. As previously shown for Gli3 (7), HA-Gli2 also binds the Gli-inhibiting protein Suppressor of Fused (SuFu), but the binding appears independent of Hh pathway activity (Fig. 2A).

**Fig.2.**
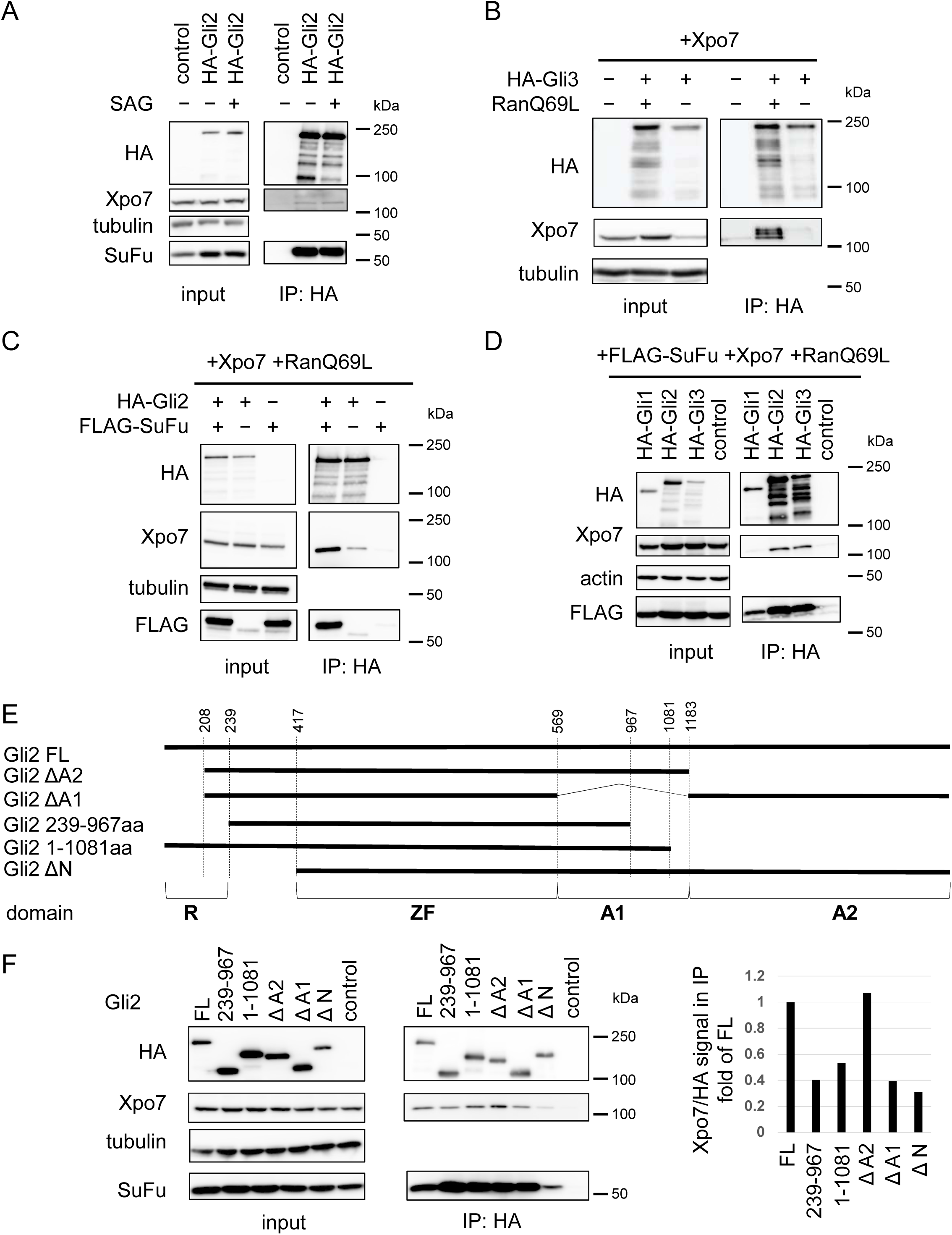
Xpo7 interacts with Gli proteins via multiple domains. **A)** NIH/3T3 cells stably expressing HA-Gli2 were left untreated or were treated with SAG, and proteins were immunoprecipitated using anti-HA-coupled beads. **B)** HA-Gli3 was co-expressed with Xpo7, with or without RanQ69L in HEK 293T cells and proteins were immunoprecipitated as in (A). The binding between Xpo7 and Gli2 is strongly enhanced in the presence of RanQ69L. **C)** HA-Gli2 was co-expressed with Xpo7, RanQ69L, with or without FLAG-SuFu in HEK293T cells and proteins were immunoprecipitated as in (A). The binding between Xpo7 and Gli2 is strongly enhanced in the presence of SuFu. **D)** HA-tagged Gli proteins (Gli1, 2 or 3) were co-expressed with Xpo7, RanQ69L, and SuFu in HEK293T cells. Immunoprecipitation was performed as in (A). Gli2 and Gli3, but not Gli1, interact with Xpo7. **E)** Schematic representation of truncated HA-tagged Gli2 constructs. All truncated Gli2 constructs additionally have serine to threonine mutations in the 6 canonical PKA target sites (P1-6A - (15)) **F)** Co-immunoprecipitation of HA-tagged Gli2 variants with Xpo7, SuFu, and RanQ69L was performed as in (A) The ratios of Xpo7/HA band intensities in the HA IP blot were calculated for all constructs as a proxy for binding strength.

To test the dependence of Gli-Xpo7 binding on RanGTP, we overexpressed HA-Gli3 and Xpo7 at high concentrations in HEK293T cells. Under these conditions, endogenous nuclear RanGTP is insufficient to promote the association between the Xpo7 and Gli3, but we were able to restore the complex by co-expressing the GTP-locked GTPase deficient RanQ69L mutant (Fig. 2B).

SuFu is a well-established inhibitor of Gli-mediated transcription and restricts full-length Gli protein nuclear accumulation (7). Interestingly, it was recently identified as one of the proteins that bind to Xpo7 in the presence of RanGTP (19). We, therefore, hypothesized that the interaction of Gli proteins and Xpo7 may depend on SuFu. Indeed, the association between HA-Gli2 and Xpo7 in HEK293T cells is stronger when the cells also overexpress FLAG-SuFu (Fig 2C). To test if Xpo7 binding is a common feature of all 3 mammalian Gli proteins, we coexpressed HA-tagged Gli1, Gli2, or Gli3 with Xpo7, SuFu, and RanQ69L. Of the three, only Gli2 and Gli3 showed strong Xpo7 binding in the presence of SuFu and RanQ69L (Fig. 2D).

To gain better insight into which domain in Gli2 is responsible for binding with Xpo7, we prepared a set of Gli2 constructs lacking one or more domains (Fig. 2E). To promote the Hh pathway-independent nuclear import of these constructs, all of them had serine-to-alanine mutations in the 6 canonical protein kinase A phosphorylation sites (P1-6A mutants - (15)). We coexpressed these HA-tagged Gli2 variants with Xpo7, SuFu, and RanQ69L in HEK293T cells and performed co-immunoprecipitation on the lysates. All truncated Gli2 proteins were able to bind to Xpo7 to some extent, but only the ΔA2 variant bound to Xpo7 with the strength comparable to the full-length Gli2 (Fig. 2F). The binding of Gli2 ΔA1 and Gli2 ΔN to Xpo7 was the weakest, suggesting that the N-terminal domain and the A1 activator domain (26) are both required for efficient Xpo7 association.

### 3. Xpo7 reduces Gli2 nuclear accumulation

To better understand if the binding of Xpo7 to Gli2/3 affects their distribution between the nuclear and cytoplasmic fractions in cells, we used immunofluorescent imaging to check the localization of HA-Gli2 in stable NIH/3T3 Flp-In cells transfected with *Xpo7* shRNAs or a control shRNA construct targeting luciferase. The shRNA constructs were shown to reduce Xpo7 expression by western blot (Fig 3A). Xpo7 depletion increased the ratio of nuclear to cytoplasmic HA-Gli2 signal in the presence of the Smo agonist SAG (Fig. 3B and C; left panel). The constitutively active P1-6A mutant of Gli2 accumulates in the nuclei at higher levels in control cells than the wild-type protein, but its localization further increases upon *Xpo7* KD (Fig. 3C; right panel).

**Fig.3.**
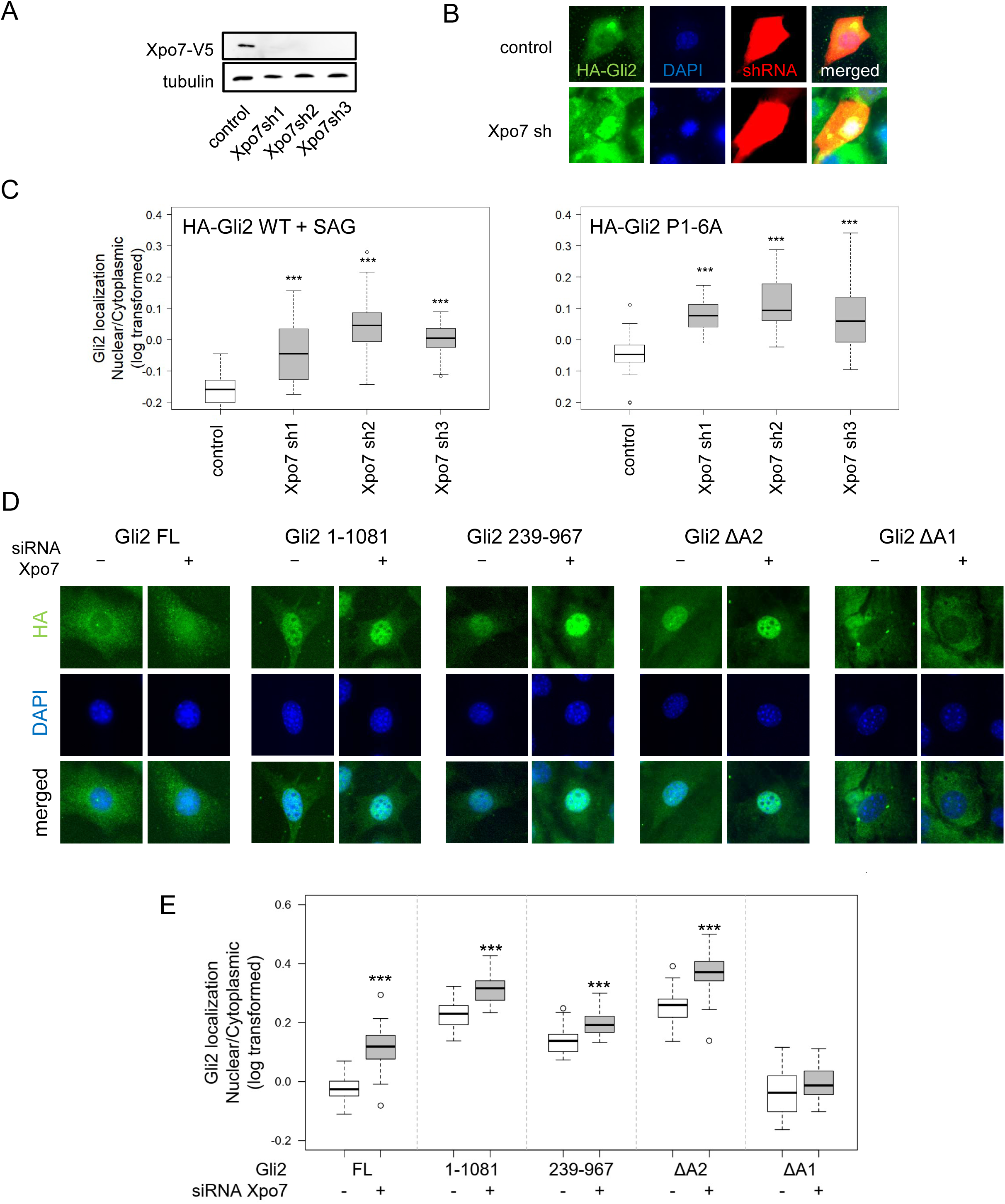
Xpo7 exports Gli2 from the nucleus. **A)** HEK293T cells co-transfected with Xpo7 and shRNAs against *Xpo7* do not have detectable levels of Xpo7 protein. **B)** Representative images of cells stably expressing HA-tagged Gli2 co-transfected with a plasmid encoding tdTomato and shRNA targeting *Xpo7* or a non-targeting control shRNA. TdTomato fluorescence (red) was used to identify shRNA-transfected cells. Cells were stained with anti-HA antibodies, counterstained with DAPI and imaged. **C)** Quantification of Gli2 localization in cells with *Xpo7* knockdown. Cells stably expressing HA-Gli2 (wild type, WT) or the constitutively active HA-Gli2 P1-6A were transiently transfected with shRNAs targeting *Xpo7* or a non-targeting control shRNA. Cells were stained as in (A). Fluorescence signal was measured in the nucleus (stained with DAPI) and cytoplasm for 30 cells per cell line and the ratio of nuclear to cytoplasmic (N/C) fluorescence was calculated for each cell. Data is shown as log_10_(N/C); n=30). **D)** Representative images of the cells stably expressing the indicated HA-tagged truncated Gli2 constructs with (+) or without (-) the addition siRNA targeting *Xpo7*. Cells were stained as in (A). **E)** Gli2 variant nuclear/cytoplasmic localization ratios were calculated as in (B) for 30 cells per cell line (data shown as log_10_(N/C); n=30). All constructs show statistically significant increase in nuclear localization upon *Xpo7* KD, except the ΔA1 construct.

To determine what domains of Gli2 are essential for the Xpo7-mediated inhibition of Gli2 nuclear accumulation, we knocked down *Xpo7* in stable cell NIH/3T3 Flp-In lines expressing each Gli2 construct previously used for coimmunoprecipitation experiments. We expected that Gli2 deletion constructs with low Xpo7 binding would not accumulate in the nucleus upon *Xpo7* KD. We observed an increase in nuclear concentration upon *Xpo7* KD for FL, 1-1081, 239-967 and ΔA2 Gli2 variants (Fig. 3D, E). There was no significant change in the localization of the ΔA1 variant upon *Xpo7* KD, which is consistent with the fact that Gli2 ΔA1 was characterized by the least Xpo7 binding in coimmunoprecipitation experiments. These results suggest that the main domain responsible for the physical and functional interaction between the Gli proteins and Xpo7 is the A1 activation domain.

## Disscussion

The shuttling of transcription factors across the nuclear membrane is one of the key regulatory mechanism in signaling cascades. In the Hh pathway specifically, the degree of target gene transcription is based on the fine balance between the amounts of Gli protein repressors and activators inside the nucleus.

The concentration of Gli proteins in the nucleus is determined by the equilibrium between their nuclear import and export. The nuclear import of Gli proteins has been extensively studied. Two putative NLS sequences are present in all three Gli proteins: NLS1 - a non-classical PY-NLS (amino acids 79-84 in Gli1), and NLS2 - a bipartite classical NLS (amino acids 383-401aa in Gli1) (14, 27). Both NLS1 and NLS2 are capable of binding the importin-α/β1. The mutation of the the two NLS sequences or the inhibition of importin β1 activity using siRNA or a drug importazole results in a reduction in nuclear accumulation of Gli1 and Gli2 (14, 28, 29). In addition to a putative role in nuclear import, the PY-NLS sequence also appears to play a role in the ciliary transport of Gli2, which is mediated by importin β2 (30).

In contrast to Gli nuclear import, the mechanism of their export form the nuclei is less well understood. Based on a recent functional screen for negative and positive regulators of the Hh pathway (16), we selected Xpo7 as the most likely candidate for the karyopherin that mediates the exit of Gli proteins from the nuclei. Xpo7 had long been considered as a specialized exportin with only two validated cargo proteins (18). Its only established physiological role so far has been in the enucleation of red blood cells (20). However, Xpo7 was recently shown to have a broad spectrum of substrates, and to act as both an export receptor and an import receptor, depending on the cargo (19). This suggests that further discoveries of the role of Xpo7 in physiological processes and in disease are likely. We show here that Xpo7 binds to Gli2/3 and negatively regulates Hh signal transduction by inhibiting the nuclear accumulation of full-length activator forms of Gli proteins.

The only exportin that had previously been implicated in Hh signaling was Crm1/Xpo1, which targets proteins containing the canonical leucine-rich nuclear export sequences (LR-NES) (31). Xpo1 was shown to participate in the nuclear export of Gli1 via a canonical LR-NES (14, 32). Interestingly, the same authors reported that the Gli2 nuclear export is mostly Xpo1-independent. Although mutation of a putative LR-NES in Gli2 somewhat increases Gli2 nuclear localization, the inhibition of Xpo1 by leptomycin B treatment does not affect the nuclear accumulation of either Gli2 (14) or Gli3 (7). These data are consistent with our results, suggesting that the primary exportin that counteracts the nuclear import of Gli2/3 is Xpo7, not Xpo1.

Unlike Xpo1, Xpo7 does not recognize simple linear LR-NES sequences. Xpo7-recognition domains contain basic residues inserted into folded structures that are significantly more difficult to discover (18). Consequently, we expected that Gli protein binding to Xpo7 may be dependent on dispersed signals and on protein structure, rather than on simple short linear motifs. Indeed, all truncated Gli2 variants we tested were able to bind to Xpo7 to some extent, suggesting that several binding sites in different Gli2 domains participate in the interaction. That said, coimmunoprecipitation experiments suggested significant differences in binding affinity of Xpo7 for different Gli2 domains. The deletion of the A2 domain does not impair Xpo7 binding to Gli2, but the deletion of the N-terminal domain or the A1 domain causes Xpo7 to bind more weakly. Accordingly, the Gli2 variant lacking the A1 domain does not accumulate in the nucleus upon the Xpo7 knockdown. Interestingly, the deletion of the C-terminal region of the A1 domain (aa 1082-1183) results in weaker Xpo7 binding (Fig. 2F) but surprisingly does not impair the ability of Xpo7 to shuttle the mutant protein from the nucleus (Fig. 3C, D). This suggests that binding affinity to Xpo7 to the protein of interest may on its own be a poor predictor of its functional role in the translocation of this protein into the cytoplasm.

We show that the binding of Gli proteins to Xpo7 is strongly enhanced in the presence of SuFu. SuFu is a well-established negative regulator of Hh signaling and was, like Xpo7, one of the top-scoring genes in the functional CRISPR/Cas9 screen for modulators of Hh activity (16). The mechanism of action of SuFu in the Hh pathway remains somewhat controversial. On the one hand, its binding to the N-terminal domains of Gli1/2 competes with the binding of importin β1/β2, respectively (28, 33). On the other hand, SuFu has been shown to localize in the nuclear fraction in association with Gli proteins (34) and to recruit transcriptional co-repressors such as p66β and the Sin3 HDAC complex to Gli-occupied chromatin loci (35, 36). We show here that an additional inhibitory role of SuFu may be to promote the Xpo7-dependent nuclear export of Gli2/3 when the pathway is inactive. SuFu itself was independently identified as an Xpo7 substrate (19), but whether the binding of SuFu to Xpo7/RanGTP occurs separately from Gli proteins is difficult to determine due to the very stable complexes formed by SuFu and Gli1/2/3 in cells. Importantly, SuFu binding to Gli proteins is not sufficient in itself to mediate Xpo7 association, since Gli2 ΔA1 retains strong SuFu binding, but shows weak Xpo7 binding and no dependence on Xpo7 for nuclear export. These results suggest that the interaction of the Gli/SuFu complex with Xpo7 depends on the specific structural configuration of that complex. SuFu binds to Gli proteins via multiple domains (37, 38) and only some of these binding events may be compatible with the formation of a quarternary Gli/SuFu/Xpo7/RanGTP complex.

Overall, it appears that SuFu may inhibit Gli proteins in different ways, some independent of Xpo7: it may inhibit Gli nuclear import, increase Gli nuclear export, or shift the balance of Gli cofactors on chromatin from coactivators to corepressors. Importantly, in cells exposed to near-saturating concentrations of the Hh ligand, SuFu remains as a top-scoring negative regulator of Hh signaling, but Xpo7 does not (Fig. 1A). In an alternative CRISPR/Cas9 screen performed only in cells treated with saturating doses of Shh, Xpo7 was in fact classified as a positive rather than negative Hh regulator (39). This suggests that Xpo7 may help fine-tune the Hh pathway in different directions depending on the concentration of the ligand.

Taking all these findings together, we postulate that Xpo7 is a regulator of the Hh pathway that cooperates with SuFu in excluding Gli2/3 from the nucleus and facilitating the maintenance of the “OFF” state for Hh signaling in the absence of ligand. Given that drugs targeting nuclear export pathways have recently found their way to the clinic (40), our data may be the stepping stone towards designing novel therapeutics against Gli-dependent tumors.

## Acknowledgements

This research was supported by FUGA5 grant 2016/20/S/NZ3/00213 from Polish National Science Centre to L.M.; P.N. and T.U. were funded by the OPUS 2017/27/B/NZ3/01163 grant from the National Science Centre. We thank Ian G. Macara for sharing his RanQ69L plasmids. We thank R. Rohatgi for sharing the anti-SuFu antibody. The authors declare no competing financial interests.

## Author contributions

L.M. and P.N. designed experiments; L.M., T.U., S.N., and P.N. performed experiments; L.M. and P.N. analyzed and interpreted data; L.M. and P. N. wrote the first draft of the manuscript; T.U. and S.N. edited the manuscript; and all authors approved the manuscript.

## Literature

1. Niewiadomski P, Niedziółka SM, Markiewicz Ł, Uśpieński T, Baran B, Chojnowska K. 2019. Gli Proteins: Regulation in Development and Cancer. Cells 8:147.

2. Mohler J. Requirements for hedgehog, a Segmental Polarity Gene, in Patterning Larval and Adult Cuticleof Drosophila 12.

3. Dessaud E, McMahon AP, Briscoe J. 2008. Pattern formation in the vertebrate neural tube: a sonic hedgehog morphogen-regulated transcriptional network. Development 135:2489–2503.

4. Jiang J, Hui C. 2008. Hedgehog Signaling in Development and Cancer. Dev Cell 15:801–812.

5. Gonnissen A, Isebaert S, Haustermans K. 2015. Targeting the Hedgehog signaling pathway in cancer: beyond Smoothened. Oncotarget 6.

6. Rohatgi R, Milenkovic L, Scott MP. 2007. Patched1 Regulates Hedgehog Signaling at the Primary Cilium. Science 317:372–376.

7. Humke EW, Dorn KV, Milenkovic L, Scott MP, Rohatgi R. 2010. The output of Hedgehog signaling is controlled by the dynamic association between Suppressor of Fused and the Gli proteins. Genes Dev 24:670–682.

8. Chen M-H, Wilson CW, Li Y-J, Law KKL, Lu C-S, Gacayan R, Zhang X, Hui C -c., Chuang P-T. 2009. Cilium-independent regulation of Gli protein function by Sufu in Hedgehog signaling is evolutionarily conserved. Genes Dev 23:1910–1928.

9. Imamoto N. 2000. Diversity in Nucleocytoplasmic Transport Pathways. Cell Struct Funct 25:207–216.

10. Güttler T, Görlich D. 2011. Ran-dependent nuclear export mediators: a structural perspective: RanGTPase-driven nuclear export. EMBO J 30:3457–3474.

11. Izaurralde E. 1997. The asymmetric distribution of the constituents of the Ran system is essential for transport into and out of the nucleus. EMBO J 16:6535–6547.

12. Ribbeck K, Kutay U, Paraskeva E, Görlich D. 1999. The translocation of transportin– cargo complexes through nuclear pores is independent of both Ran and energy. Curr Biol 9:47–S1.

13. Görlich D, Kutay U. 1999. Transport Between the Cell Nucleus and the Cytoplasm. Annu Rev Cell Dev Biol 15:607–660.

14. Barnfield PC, Zhang X, Thanabalasingham V, Yoshida M, Hui C. 2005. Negative regulation of Gli1 and Gli2 activator function by Suppressor of fused through multiple mechanisms. Differentiation 73:397–405.

15. Niewiadomski P, Kong JH, Ahrends R, Ma Y, Humke EW, Khan S, Teruel MN, Novitch BG, Rohatgi R. 2014. Gli protein activity is controlled by multisite phosphorylation in vertebrate Hedgehog signaling. Cell Rep 6:168–181.

16. Pusapati GV, Kong JH, Patel BB, Krishnan A, Sagner A, Kinnebrew M, Briscoe J, Aravind L, Rohatgi R. 2018. CRISPR Screens Uncover Genes that Regulate Target Cell Sensitivity to the Morphogen Sonic Hedgehog. Dev Cell 44:113-129.e8.

17. Kutay U, Hartmann E, Treichel N, Calado A, Carmo-Fonseca M, Prehn S, Kraft R, Görlich D, Bischoff FR. 2000. Identification of Two Novel RanGTP-binding Proteins Belonging to the Importin β Superfamily. J Biol Chem 275:40163–40168.

18. Mingot J-M, Bohnsack MT, Jäkle U, Görlich D. 2004. Exportin 7 defines a novel general nuclear export pathway. EMBO J 23:3227–3236.

19. Aksu M, Pleiner T, Karaca S, Kappert C, Dehne H-J, Seibel K, Urlaub H, Bohnsack MT, Görlich D. 2018. Xpo7 is a broad-spectrum exportin and a nuclear import receptor. J Cell Biol 217:2329–2340.

20. Hattangadi SM, Martinez-Morilla S, Patterson HC, Shi J, Burke K, Avila-Figueroa A, Venkatesan S, Wang J, Paulsen K, Görlich D, Murata-Hori M, Lodish HF. 2014. Histones to the cytosol: exportin 7 is essential for normal terminal erythroid nuclear maturation. Blood 124:1931–1940.

21. Dorfman J, Macara IG. 2008. STRAD␣ Regulates LKB1 Localization by Blocking Access to Importin-␣, and by Association with Crm1 and Exportin-7. Mol Biol Cell 19:13.

22. Niewiadomski P, Rohatgi R. 2015. Rapid Screening of Gli2/3 Mutants Using the Flp-In System. Methods Mol Biol Clifton NJ 1322:125–130.

23. Ran FA, Hsu PD, Wright J, Agarwala V, Scott DA, Zhang F. 2013. Genome engineering using the CRISPR-Cas9 system. Nat Protoc 8:2281–2308.

24. Gonçalves C, Gross F, Guégan P, Cheradame H, Midou P. 2014. A robust transfection reagent for the transfection of CHO and HEK293 cells and production of recombinant proteins and lentiviral particles - PTG1. Biotechnol J 9:1380–1388.

25. Sasaki H, Hui C, Nakafuku M, Kondoh H. A binding site for Gli proteins is essential for HNF-3β floor plate enhancer activity in transgenics and can respond to Shh in vitro 10.

26. Sasaki H, Nishizaki Y, Hui C, Nakafuku M, Kondoh H. 1999. Regulation of Gli2 and Gli3 activities by an amino-terminal repression domain: implication of Gli2 and Gli3 as primary mediators of Shh signaling. Dev Camb Engl 126:3915–3924.

27. Hatayama M, Aruga J. 2012. Gli Protein Nuclear Localization Signal, p. 73–89. *In* Vitamins & Hormones. Elsevier.

28. Szczepny A, Wagstaff KM, Dias M, Gajewska K, Wang C, Davies RG, Kaur G, Ly-Huynh J, Loveland KL, Jans DA. 2014. Overlapping binding sites for importin β1 and suppressor of fused (SuFu) on glioma-associated oncogene homologue 1 (Gli1) regulate its nuclear localization. Biochem J 461:469–476.

29. Torrado B, Graña M, Badano JL, Irigoín F. 2016. Ciliary Entry of the Hedgehog Transcriptional Activator Gli2 Is Mediated by the Nuclear Import Machinery but Differs from Nuclear Transport in Being Imp-α/β1-Independent. PLOS ONE 11:e0162033.

30. Han Y, Xiong Y, Shi X, Wu J, Zhao Y, Jiang J. 2017. Regulation of Gli ciliary localization and Hedgehog signaling by the PY-NLS/karyopherin-β2 nuclear import system. PLoS Biol 15.

31. Fornerod M, Ohno M, Yoshida M, Mattaj IW. 1997. CRM1 is an export receptor for leucine-rich nuclear export signals. Cell 90:1051–1060.

32. Kogerman P, Grimm T, Kogerman L, Krause D, Undén AB, Sandstedt B, Toftgård R, Zaphiropoulos PG. 1999. Mammalian Suppressor-of-Fused modulates nuclear– cytoplasmic shuttling of GLI-1. Nat Cell Biol 1:312–319.

33. Shi Q, Han Y, Jiang J. 2014. Suppressor of fused impedes Ci/Gli nuclear import by opposing Trn/Kap 2 in Hedgehog signaling. J Cell Sci 127:1092–1103.

34. Zhang Z, Shen L, Law K, Zhang Z, Liu X, Hua H, Li S, Huang H, Yue S, Hui C, Cheng SY. 2017. Suppressor of Fused Chaperones Gli Proteins To Generate Transcriptional Responses to Sonic Hedgehog Signaling. Mol Cell Biol 37:e00421-16, /mcb/37/3/e00421-16.atom.

35. Lin C, Yao E, Wang K, Nozawa Y, Shimizu H, Johnson JR, Chen J-N, Krogan NJ, Chuang P-T. 2014. Regulation of Sufu activity by p66β and Mycbp provides new insight into vertebrate Hedgehog signaling. Genes Dev 28:2547–2563.

36. Cheng SY, Bishop JM. 2002. Suppressor of Fused represses Gli-mediated transcription by recruiting the SAP18-mSin3 corepressor complex. Proc Natl Acad Sci 99:5442–5447.

37. Oh S, Kato M, Zhang C, Guo Y, Beachy PA. 2015. A Comparison of Ci/Gli Activity as Regulated by Sufu in Drosophila and Mammalian Hedgehog Response. PloS One 10:e0135804.

38. Merchant M, Vajdos FF, Ultsch M, Maun HR, Wendt U, Cannon J, Desmarais W, Lazarus RA, de Vos AM, de Sauvage FJ. 2004. Suppressor of Fused Regulates Gli Activity through a Dual Binding Mechanism. Mol Cell Biol 24:8627–8641.

39. Breslow DK, Hoogendoorn S, Kopp AR, Morgens DW, Vu BK, Kennedy MC, Han K, Li A, Hess GT, Bassik MC, Chen JK, Nachury MV. 2018. A CRISPR-based screen for Hedgehog signaling provides insights into ciliary function and ciliopathies. Nat Genet 50:460–471.

40. Wang AY, Liu H. 2019. The past, present, and future of CRM1/XPO1 inhibitors. Stem Cell Investig 6.

